# Differences in sustained attention but not distraction in preschoolers from lower socioeconomic status backgrounds

**DOI:** 10.1101/2021.04.06.438161

**Authors:** Roxane S. Hoyer, Eric Pakulak, Aurélie Bidet-Caulet, Christina M. Karns

## Abstract

In children, the ability to listen to relevant auditory information and suppress distracting information is a foundational skill for learning and educational achievement. Distractibility is supported by multiple cognitive components (voluntary attention orienting, sustained attention, distraction, phasic arousal, as well as impulsivity and motor control) that may mature at different ages. Here we used the Competitive Attention Test (CAT) to measure these components in 71 4- and 5-year-old children. The goal of this study was to characterize the changes in efficiency of attention during the preschool period, and to explore differences in distractibility in preschool children that could be related to the socioeconomic status (SES) background of the child’s family. We found that sustained attention improves from age 4 to 5, while voluntary attention orienting is still immature during the preschool period. In addition, independent of age, task-irrelevant sounds induced distraction, phasic arousal, and impulsivity. Children from lower SES backgrounds showed reduced sustained attention abilities and increased impulsivity. However, 3-year-old children and a minority of 4- and 5-year-olds did not manage to perform the task according to the instructions; the CAT thus seems suitable to assess distractibility only in preschoolers with sufficiently developed sustained attention skills to efficiently complete the task. Taken together, the present findings suggest that distractibility is still developing during the preschool period and is likely to vary depending on the SES background of a child’s family.

## Introduction

In children, the ability to listen to relevant auditory information and suppress distracting information is a foundational skill for learning and educational achievement (Stevens & Bavelier, 2012). For example, in a typical classroom, a child needs to listen to a teacher’s instructions or to focus on a task when there are distractors in the environment, such as other children talking. The common term “paying attention” implies that attention is a unitary phenomenon, but it is in fact a multifaceted construct supported by multiple brain networks that undergo significant and differential development in childhood (Posner et al., 2014; Wetzel & Schröger, 2014). The present study focuses on the different facets of distractibility by examining differences in behavioral performance during an attention task in 4- and 5-year-old children from different familial socioeconomic status (SES) backgrounds.

### Distractibility

The term ‘distractibility’ has been used to describe an attention state that determines the propensity to have attention captured by irrelevant information and to react behaviorally to this information (Bidet-Caulet et al., 2015; ElShafei et al., 2019, 2020; Hoyer et al., 2021; Masson & Bidet-Caulet, 2019). Distractibility is a result of the balance between voluntary and involuntary attention processes.

Voluntary attention is goal-directed and can enhance the processing of relevant features (Corbetta & Shulman, 2002), locations (Posner, 1980), or modalities (Karns et al. 2009). Two forms of voluntary attention directly contribute to distractibility: voluntary orienting and sustained attention. First, voluntary orienting allows one to selectively focus on a location and engenders perceptual enhancement of upcoming targets occurring at this location (Petersen & Posner, 2012; Posner, 1980; Yantis & Jonides, 1990). Behaviorally, voluntary orienting can be measured by contrasting reaction time (RT) to targets preceded by uninformative and informative cues (Petersen & Posner, 2012; Posner, 1980). Second, sustained attention is the ability to maintain attentional focus over time on a given task, and relies strongly on tonic arousal, also called vigilance (Betts et al., 2006; Oken et al., 2006; Parasuraman et al., 1989). Sustained attention can vary over multiple time scales, from rapid (across milliseconds) to slow (e.g., over a day), with attentional fluctuations over time (for review, see Esterman & Rothlein, 2019). Thus, voluntary orienting and sustained attention allow enhanced processing of ongoing task-related relevant stimuli and the effective maintenance of this enhancement over time. Overall, these two forms of voluntary attention allow one to focus.

In contrast to voluntary attention, involuntary attention can be oriented toward unexpected salient stimuli, leading to distraction when the stimulus is irrelevant in the task context (Bidet-Caulet et al., 2015; Escera, Alho, et al., 2000; Näätänen, 1992). “Distraction,” distinct from distractibility, describes the deleterious impact of a distractor on ongoing cognitive and behavioral performance. Indeed, when unexpected irrelevant salient stimuli occur in a task context, the reactive allocation of attention and resources toward the irrelevant event is followed by a reallocation of resources toward the task: RT and accuracy can thus, respectively, increase and decrease as a result of distraction (Bidet-Caulet et al., 2015; Escera, Alho, et al., 2000; Näätänen, 1992). In this respect, distractors occurring while one is processing relevant information usually disturb the processing of this information and have a deleterious impact on behavioral performance.

Counterintuitively, a distractor that occurs suddenly in the environment can also have a facilitation effect on RT. Since attention and arousal (i.e., alertness induced by the noradrenergic system) networks are interconnected in the brain (Aston-Jones & Cohen, 2005), this facilitation effect is induced by a transient increase in noradrenaline in response to the occurrence of a distractor and leads one to temporarily react more quickly to any upcoming stimuli (Bidet-Caulet et al., 2015; Masson & Bidet-Caulet, 2019; Max et al., 2015; Näätänen, 1992; Wetzel et al., 2012, SanMiguel et al., 2010). Interestingly, an increased false alarm rate is typically observed in impulsive people and might ensue from an enhanced phasic arousal effect (Eysenck & Eysenck, 1985; Houston & Stanford, 2001; Zhang et al., 2015) and reduced motor control (Booth et al., 2003; Kanaka et al., 2008; van den Wildenberg & Crone, 2005; Wright et al., 2003).

As multiple cognitive components (voluntary attention orienting, sustained attention, distraction, phasic arousal, as well as impulsivity and motor control) contribute to distractibility, it is fair to assume that distractibility is not a unitary function, but rather a cognitive state.

### Distractibility in preschool aged children

Evidence suggests that children are more distractible than adults (Hoyer et al., 2021; Wetzel et al., 2006, 2016; Wetzel & Schröger, 2014). However, there remains a gap in our understanding of the origins of immature distractibility during early childhood. Below we discuss results from previous studies that separately investigated the different components of distractibility in preschool-aged children.

#### Voluntary attention orienting and sustained attention

Studies using tasks with endogenous cues that are either informative or uninformative have yielded conflicting results on the efficacy of voluntary attention in preschool children: some studies suggest that the capacity to voluntarily orient attention reaches an adult-like efficiency before the age of four (Colombo, 2001; Johnson et al., 1991; Ross-Sheehy et al., 2015) while others show that the benefit in RT to targets following informative cues increases from four to six years of age (Hrabok et al., 2007; Mezzacappa, 2004; Posner et al., 2014; Rothbart et al., 2011). In preschool-aged children, sustained attention has been investigated using various child-friendly tasks. Taken together, results suggest that sustained attention emerges within the first years of life (Kanaka et al., 2008; Reynolds & Romano, 2016) and these skills have been found to progressively improve between three and five years of age (Graziano et al., 2011; Reynolds & Romano, 2016; Richards & Casey, 1991; Ruff & Capozzoli, 2003). Behaviorally, these changes in sustained attention are reflected in a gradual reduction of false alarm and missed response rates, as well as in RT variability (Mahone et al., 2001). To date, few studies have investigated the impact of isolated distracting events on sustained attention abilities; the small number of available studies suggests that efficient sustained attention abilities in children may shield against the deleterious effect of distraction (Oakes & Tellinghuisen, 1994; Slobodin et al., 2015).

#### Distraction

To our knowledge, only one study has investigated the distraction effect on behavioral performance in preschool-aged children. In an audio-visual oddball task, in which participants need to discriminate targets preceded by irrelevant standard, deviant, or novel sounds, the deleterious effect of distraction - namely, increased reaction times to targets preceded by novel or deviant compared to standard sounds – was found to progressively decrease during the preschool period, with a critical developmental step between ages four and five (Wetzel et al., 2018). However, it remains to be determined whether this difference between four and five-year-old children can also be observed when distractors are not deviant or novel and not embedded in a stream of standard sounds, as it is the case in oddball paradigms.

#### Phasic arousal

To date, no study has investigated developmental differences, during the preschool years, in the beneficial behavioral effect which can be induced by distracting sounds. Phasic arousal, which is hypothesized to be responsible for increased speed in RT to targets preceded by a distractor, has been studied mostly using physiological measurements in the first year of life (Hernes et al., 2002). For example, an increased pupil dilation response to rare, unexpected, and complex sounds has been observed in infants from 13 to 16 months of age compared to adults (Wetzel et al., 2015; Max et al., 2015), suggesting that phasic arousal is increased in infants. Behaviorally, phasic arousal has also been found to be increased in children aged 6 to 13-years-old compared to adults (Hoyer et al., 2021). To that extent, it is possible that phasic arousal undergoes developmental changes during the preschool period; to our knowledge, no studies have yet attempted to answer this question.

#### Motor control and impulsivity

Sensory and motor areas of the brain are typically the first to mature (Casey et al., 2005) with structural changes in the sensorimotor cortex leading to an adult-like functioning between late infancy and the preschool period. However, motor control is dependent on many interconnections between cortical and sub-cortical regions of the brain (e.g., the prefrontal and lateral temporal cortices) which do not appear to reach a complete level of maturity until young adulthood (Gogtay et al., 2004). To that extent, motor control and impulsivity are likely to influence the motor response, which can in turn lead young children to react randomly.

Studies of the development of the different components of distractibility during the preschool years suggest that sustained attention seems to increase during this period, but report contradictory results regarding voluntary orienting. Furthermore, the studies that have examined distraction, phasic arousal, as well as motor control and impulsivity are too few to draw strong conclusions about the behavioral differences associated with the development of these mechanisms during the preschool period. Thus, to better understand how distractibility develops during this period, it is necessary to simultaneously assess its components in children.

### The Competitive Attention Test

The Competitive Attention Test (CAT) is a detection paradigm designed to assess the behavioral and brain correlates of distractibility (Bidet-Caulet et al., 2015). The advantage of this paradigm is that it combines the Posner attention-network task and principles of the oddball paradigm to provide simultaneous and dissociated measures of voluntary attention, distraction, phasic arousal, motor control and impulsivity in children and adults (Hoyer et al., 2021). To assess voluntary attention orienting, the CAT includes informative and uninformative visual cues toward the spatial location of a forthcoming auditory target. To measure distraction, the CAT includes trials with a task-irrelevant distracting sound preceding the target at several different delays. This change in distractor timing onset allows the dissociation of the behavioral effects of facilitating phasic arousal (the difference between median RT in no-distractor and early-distractor conditions) and detrimental distraction (the difference between median RT in late- and early-distractor conditions). Based on previous results (Bidet-Caulet et al., 2015; Masson & Bidet-Caulet, 2019), these differences can be interpreted respectively as good approximations of the facilitation and detrimental distraction effects triggered by so-called distracting sounds (see Appendix A Fig. A1). Results from studies in adults typically show that the voluntary orienting effect is manifested by a RT reduction in informative compared to uninformative trials; the distraction effect is manifested by increased RT in late compared to early-distractor condition; and, finally, the phasic arousal effect is indexed by a RT reduction in the early distractor condition compared to no distractor condition.

In order to study the development of these facets of attention with more precision, the CAT was recently adapted for young children and used in a study of a large cohort of participants aged 6 to 25 years (Hoyer et al., 2021). In this study, the behavioral measurement parameters of the CAT were refined compared to those previously used in adults (see Table 2 and Method for a detailed description). Results showed that voluntary orienting reaches an adult-like efficacy at 6 years of age, while the ability to sustain attention gradually develops from 8 to 12 years of age; interestingly, distraction is manifested as omissions (i.e., missed targets) of relevant stimuli in 6-7-year-olds and as impulsivity (i.e., responses to distractors) in 11-12-year-olds. However, the RT distraction measure was not modulated by age, while the RT facilitation effect linked to phasic arousal decreased from 6 to 12 years of age. This attentional imbalance, resulting in increased distractibility in children, may then be more related to reduced sustained attention capacities, enhanced distraction, and increased arousal effects in childhood (6-8-year-olds), but to increased impulsivity in older children and adolescents (10-17-year-olds). Importantly, some measurements (e.g., missed responses, RT variability) show higher variability in younger children than in adults. A part of this variability may be related to environmental factors such as socioeconomic status.

### Socioeconomic Status

Socioeconomic status (SES) background is a proxy variable for variability in the early environment, typically assessed by a combination of parental education, occupational status, and/or household income. Disparities as a function of SES have been documented in a wide range of neurocognitive outcomes, and one of the neurocognitive systems most consistently associated with SES is self-regulation, including specific aspects of attention (Hackman et al., 2010; Pakulak et al., 2018; Ursache & Noble, 2016). Altered functioning of executive attention has been linked to reduced voluntary attention abilities (Diamond, 2013; Posner, 1980, 2012), and has been consistently reported in children and adolescents from lower SES (LSES) backgrounds (Farah, 2017; Farah et al., 2006; Noble et al., 2013). Event-related brain potential studies have also found differences in selective attention as a function of SES in adolescents (D’Angiulli et al., 2008) and in preschool-aged children (Giuliano et al., 2018; Hampton Wray et al., 2017; Stevens et al., 2009). These studies have found that children from LSES backgrounds show an increased brain response to stimuli they are instructed to ignore, relative to children from higher SES (HSES) backgrounds. Moreover, the relationship between SES and brain response to distracting sounds is mediated by the sympathetic nervous system activity: the larger the sympathetic activity the better distractor suppression, suggesting a biological cost, for children from LSES backgrounds, to achieve similar cognitive performance to children from HSES backgrounds (Giuliano et al., 2018).

Increasingly, evidence suggests that the effects of socioeconomic inequality begin early and persist during development. For example, behavioral signs of impulsivity in the first year of life persist into first grade only in children from LSES backgrounds (Meade, 1981); in this population, impulsivity in adolescence has also been linked to increased risk taking in early adulthood (Auger et al., 2010). Importantly, this pattern of results may represent a functional adaptation to environmental demands (e.g., increased sensitivity to potential threats) that may have deleterious effects in other environments (e.g., a classroom). In addition, these systems are amenable to evidence-based interventions targeting self-regulation and attention in preschool-aged children (Neville et al., 2013). In order to inform the refinement of such approaches and the development of novel approaches, it is crucial to improve our understanding of specific aspects of attention that contribute to distractibility in preschool-aged children from different SES backgrounds.

### The Current Study

As the development of distractibility during the preschool years is still an open question, the goals of the present exploratory study are (i) to characterize differences in efficiency of multiple aspects of attention during the preschool period, and (ii) to explore differences in distractibility in preschool-aged children that could be related to the SES background of the child’s family. Specifically, we examine the development of voluntary orienting and sustained attention, distraction, phasic arousal, impulsivity and motor control, as well as potential differences in this development as a function of SES. Based on the literature, we hypothesized that distraction would decrease during early childhood, accompanied by an improvement in motor control and by a maturation of voluntary orienting and sustained attention. We also expected to observe a different profile in children from LSES backgrounds, with reduced voluntary attention (voluntary orienting and sustained attention) and increase distraction and impulsivity compared to their peers from HSES backgrounds.

## Methods

### Participants

A total of one hundred and six preschool children aged three (3YO) four (4YO) or five (5YO) years participated at their usual childcare site (sample before exclusion, see below and Table 1 for the final sample). All children had normal or corrected-to-normal vision. Parents were informed and provided signed informed consent on-site, and children provided informed verbal assent prior to participation. With their permission, parents at each site were entered to win a $50 gift card in a lottery regardless of whether they opted for their child to participate in the study. Children chose a small educational prize for participating in the study. Recruitment and study procedures were approved by the University of [blinded] Research Compliance Services and by participating preschools.

**Table 1.** Characteristics of the population sample.

|  | Sample size |  | Age in months | Gender |  |
| --- | --- | --- | --- | --- | --- |
|  | *Included* | *excluded* |  | *Female* | *Male* |
| **Total** | 71 | 21 | 60.0 ± 0.7 | 53.8% | 46.2% |
| **Lower SES** | 43 | 16 | 60.7 ± 0.9 | 55.8% | 44.2% |
| **Higher SES** | 28 | 5 | 58.9 ± 1.2 | 42.9% | 57.1% |

*Note.* Detailed samples by SES for age in months (± SEM) and gender.

Socioeconomic status was operationalized based on demographic characteristics of participating preschool sites, which were selected specifically for this purpose. While individual-level assessment of SES was beyond the scope of this study, parental income and education are considered a valuable proxy for the wider range of characteristics that tend to vary as a function of SES, even at an aggregate group level (Hackman et al., 2010; Ursache & Noble, 2016). Children were recruited from either preschools associated with a university campus (higher SES; HSES) or from Head Start preschool sites (lower SES; LSES). Note that, for readability, we will use “HSES and LSES children” in the Method and Results sections of this manuscript to refer to “children from higher SES and lower SES backgrounds” respectively.

Head Start preschools (LSES preschool sites) are part of a U.S. program serving families living at or below the poverty line, which is determined by the U.S. Federal Poverty Guidelines (https://aspe.hhs.gov/poverty-guidelines); e.g., for the year in which data were acquired the poverty line for a family of four was an annual salary of $25,100 (see Appendix B Table B1 for guidelines for different family sizes). University sites (HSES preschool sites) serve children whose parents are faculty, students, or staff at the university. According to the information provided by the university sites, among children from the HSES preschool sites, 60% had parents who were faculty members, 11% had parents who were students at the university and 29% had parents who were staff at the university (e.g., administrative staff). Among them, less than 5% could benefit from the federal “reduced lunch program”, for which eligibility criteria are considerably more strict than poverty guidelines, e.g., for the year in which data were acquired the criteria for a family of four was an annual salary of $46,435 (see Appendix B Table B1 for guidelines for different family sizes). While this operationalization was necessarily broad and imprecise, it is likely that the two groups were qualitatively different in ways that reflect to some degree differences captured by more systematic assessments of SES.

A total of ninety-two 4-5YO children participated in the study, but data from 21 children were excluded due to non-compliance with instructions (e.g., not looking at the screen, not sitting still, talking or singing during the test): specifically, 16 LSES and 5 HSES children were excluded. We initially planned to include 3YO children. After testing 14 3YO children and evaluating their data, we decided to cease testing 3YO because they had difficulty maintaining focus on the task. They had an average of 40% of correct responses and a high rate of missed responses (32%) or other errors (cue responses: 4%, random responses: 5%, distractor responses: 4%, anticipated responses: 10%, and late responses: 5%). Thus, data from 3YO children were not included in the analysis, leaving a final sample of 71 subjects (43 LSES and 28 HSES, 54% female, 4 to 5 years of age). Table 1 summarizes characteristics of the final sample.

### Stimuli and task

Detailed task methods have been previously published (see Hoyer et al., 2021 and Appendix C for more details). In brief, children were instructed to keep their eyes fixated on a cross during the task intervals between trial events. In informative trials (75%), a visual cue (a dog facing left or right, 200 ms duration) indicated in which ear (left or right, respectively) the target sound (a dog bark, Fig. 1a) would be played (200 ms duration) after a delay (940 ms). Uninformative trials (25%) were constructed identically, but the cue was a dog facing straight ahead. The target sound was monaurally presented in headphones. For 50% of trials, an additional 300-ms duration binaural distracting sound (18 different possible ringing sounds) was played during the 940 ms delay period (Fig. 1b). The distracting sound could be played at three different times during the delay: 200 ms (Dis1), 400 ms (Dis2) and 600 ms (Dis3) after cue offset, distributed equiprobably. Target sounds were played at 75 dBA and distracting sounds at 85 dBA. Cue categories (informative, uninformative) and target categories were equiprobable for trials with and without distracting sounds (no distractor: NoDis, distractor: Dis). Positive feedback (happy dog holding a bone in his mouth) was presented when participants detected the target within 3300 ms after onset (800-ms duration, 500 ms after the response), followed by a fixation period (1700 ms to 1900 ms). If the participant did not respond in time, the fixation cross was displayed for an additional randomized delay (0 ms to 200 ms). Participants were asked to press the same key as fast as possible when they heard the left or right target sound, and were asked to focus their attention to the cued side.

**Figure 1.**
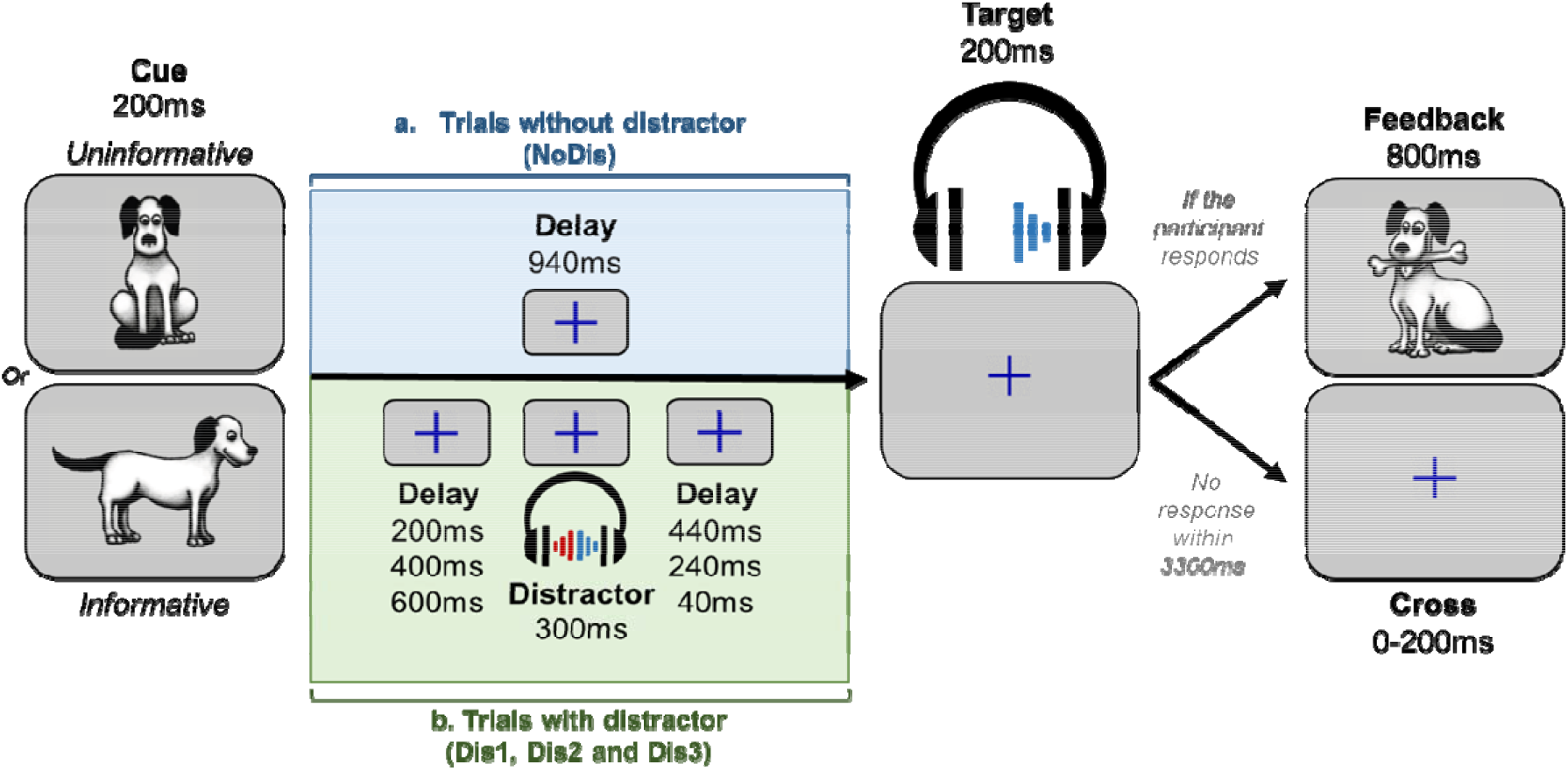
Protocol. *Note*. a) In uninformative trials, a front-facing dog was used as visual cue (informative or uninformative), indicating that the target sound would be played in either the left or right ear after a delay. If the participant gave a correct answer, a feedback cue was displayed. b) In trials with distractors the task was similar, but a binaural distracting sound - such as a phone ring - was played during the delay between cue and target. The distracting sound could equiprobably onset at three different times: 200 ms, 400 ms, or 600 ms after the cue offset.

### Procedure

Participants were tested in groups of two in a quiet room. The composition of each duo was determined by the classroom teachers as two children who were considered to have good interactions at school but did not regularly play together during playtimes. This was done to minimize both potential stress (children were all tested by two testers they only had met once during the presentation of the study to the class, so we reasoned that being with a classmate during the experiment would make the situation more comfortable for children) and distraction (children are often tempted to talk to close friends even when asked to not do so) during the experimental session. Prior to the task, children were shown a treasure map (see Fig. 2). At the end of each block, children took a break to sing a nursery song with experimenters while pretending to row to the next island. If children chose not to complete all three blocks, they still received a prize.

**Figure 2.**
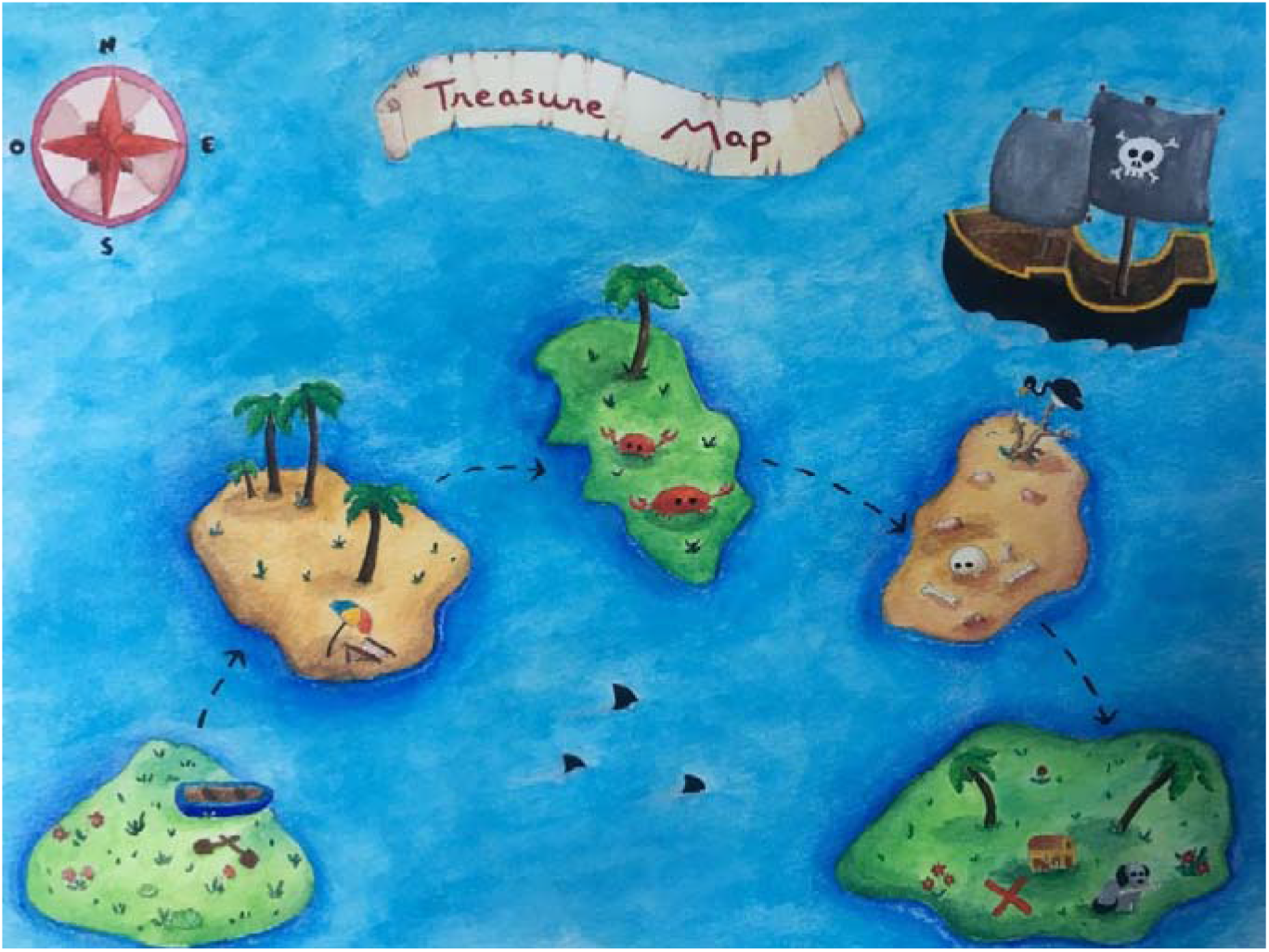
Treasure map used to motivate children during testing. In each block they would help a dog find his bone by pressing the button when they heard the dog bark. Children were told to press the button with their dominant hand (the hand they preferentially use to draw) only when they heard the dog bark, and to ignore distracting sounds. Whatever the direction of the cue, experimenters explained to children that they always needed to press the same button (the only response button - the left touchpad key - marked with a dog sticker to answer). Additional visual illustrations were used by the experimenters to make sure that children understood the instructions (see Fig. 3). To ensure that children understood the difference between the left and right (direction of the cue, informative cue), experimenters used a printed illustration of the left and right cue and asked to each child to point the ear in which the dog was going to bark. All children were able to appropriately indicate in which ear the dog was expected to bark following the left or right cue presentation. Experimenters also made sure that all the children were able to explain that the forward-facing dog (uninformative cue) meant that they could not know in which ear the target sound would be played.

**Figure 3.**
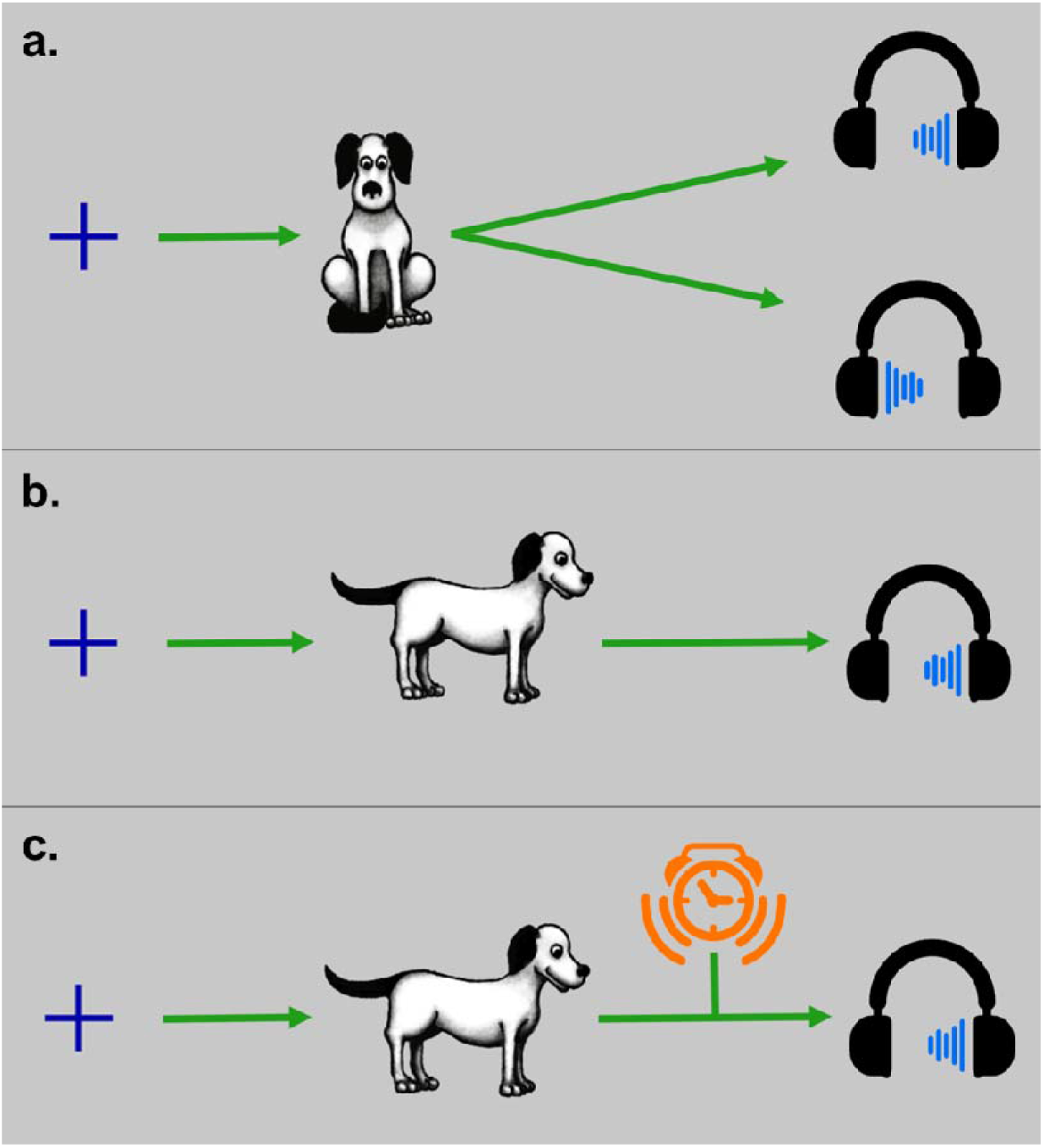
Additional illustrations for instructions. *Note*. a) Depiction of an uninformative NoDis trial, b) an informative NoDis trial and c) a distractor trial.

During the task, participants were seated in front of a laptop (approximately 50 cm from the screen) that presented pictures and sounds and recorded responses using Presentation software (Neurobehavioral Systems, Albany, CA, USA). Auditory stimuli were played in headphones. Participants performed three 4-minutes blocks of 48 pseudo-randomized trials each. Each child verbally confirmed to the experimenter that they were able to hear the dog bark during the task. An experimental session lasted around 30 minutes.

### Measurement parameters

We used a custom MATLAB program to preprocess data. The shortest RT for a correct response (RT lower limit) was calculated for each age range (see Appendix D Fig. D1). For each participant, the longest RT for a correct response (RT upper limit) was calculated from all RT > 0 ms using the Tukey method of leveraging the interquartile range. Correct response rate corresponds to the percentage of responses with a RT (relative to target onset) superior or equal to RT lower limit and inferior or equal to RT upper limit. Eight behavioral measures were extracted for each child (Table 2, see also Appendix E Fig. E1).

**Table 2.** Response types, parameters for computations and interpretation of responses.

| Behavioral measurements of attention effects on target processing |  |  |
| --- | --- | --- |
| Response Type | Description | Response Interpretation |
| Reaction times | RT of positive response times | Overall processing speed |
| Arousal effect | Median RT: NoDis – Dis1 | Facilitation effect:<br>Index of arousal due to presence of early distractor |
| Distraction effect | Median RT: Dis3 – Dis1 | Distraction effect:<br>Index of competition between distractor and target |
| Standard deviation of reaction times | Mean standard deviation of positive response times in the NoDis condition for each block separately | Variability in processing speed<br>Sustained attention |
| Late response | Percentage of responses in the NoDis condition during the period starting from the RT upper-limit to 3300 ms | Slow processing error<br>Failure of short term sustained attention |
| Missed response | Percentage of trials without any response made during the entire trial duration up to 3300 ms post-target | Error of omission<br>Lapse of sustained attention |
| Behavioral measurements of attention effects on target expectancy |  |  |
| Response Type | Description | Response Interpretation |
| Cue response | Percentage of [incorrect or false-alarm] responses performed during the 150-450ms period post-cue onset | Erroneous response to the cue<br>Impulsivity |
| Distractor response | Percentage of responses performed during the 150-450 ms period post-distractor onset | Erroneous response to the distractor<br>Impulsivity |
| Anticipated response | Percentage of responses performed:<br>In NoDis and Dis1: from 300 ms pre-target to the RT lower limit post-target<br>in Dis2: from 150 ms pre-target to the RT lower limit post-target<br>in Dis3: from 100 ms post-target to the RT lower limit post-target | Erroneous response in anticipation of the target<br>Impulsivity |
| Random responses | Percentage of responses performed in the remaining periods of the trials:<br>in NoDis: during the 450 to 850 ms period post-cue onset<br>in Dis1: during the 450 to 550 ms period post-cue onset<br>in Dis2: during the 450 to 750 ms period post-cue onset<br>in Dis3: during the 450 to 950 ms period post-cue onset | Erroneous responses outside of response parameters above |

### Statistical analysis

#### Data analysis

We expected large inter-individual variability in RT and response type rates as a function of condition, which limits the comparison of data between conditions and means that data cannot simply be pooled for analysis. Generalized Linear Mixed Models (GLMM) are preferred, as they allow for correction of systematic variability (Bates et al., 2015). The heterogeneity of performance between subjects and experimental conditions was considered by defining them as effects with random intercepts and slopes, thus instructing the model to correct for any systematic differences in variability between the subjects (between-individual variability). The marginal (variance explained by fixed factors) and conditional (variance explained by fixed and random factors) R^2^ for each models have been computed using the Nakagawa & Schielzeth’s method (2013), R MuMIn package, and can be found in Table 4.

To assess the impact of the manipulated task parameters (cue information and distractor type) and participant demographic characteristics (age in months and SES), on each type of behavioral measure (RT, RT standard deviation, late responses, missed responses, cue responses, distractor responses, anticipated responses, random responses), we analyzed the influence of four possible fixed effects (unless specified in Table 2): the between-subject factor AGE: continuous (min: 48, max: 71); the between-subject factor SES: 2 levels (LSES and HSES); the within-subject factor CUE: 2 levels (CUE: informative vs. uninformative); the within-subject factor DISTRACTOR: 4 levels (NoDis, Dis1, Dis2 and Dis3), except for distractor responses: 3 levels (Dis1, Dis2 and Dis3). The Block factor: 3 levels (1^st^, 2^nd^ and 3^rd^), was used in the RT standard deviation analysis to investigate sustained attention along the task. A summary of the data and factors used in statistical modeling can be found in Table 3. Note that for response types cumulating less than a mean of 10 observations across subjects (cue responses, distractor responses and late responses), we did not consider the within-subject factor DISTRACTOR in the analysis. Although the age (in months) was considered in all models, we did not model this variable in interaction with others to avoid the detection of a misleading AGE*SES interaction, which could potentially be driven by small sample sizes.

**Table 3.**
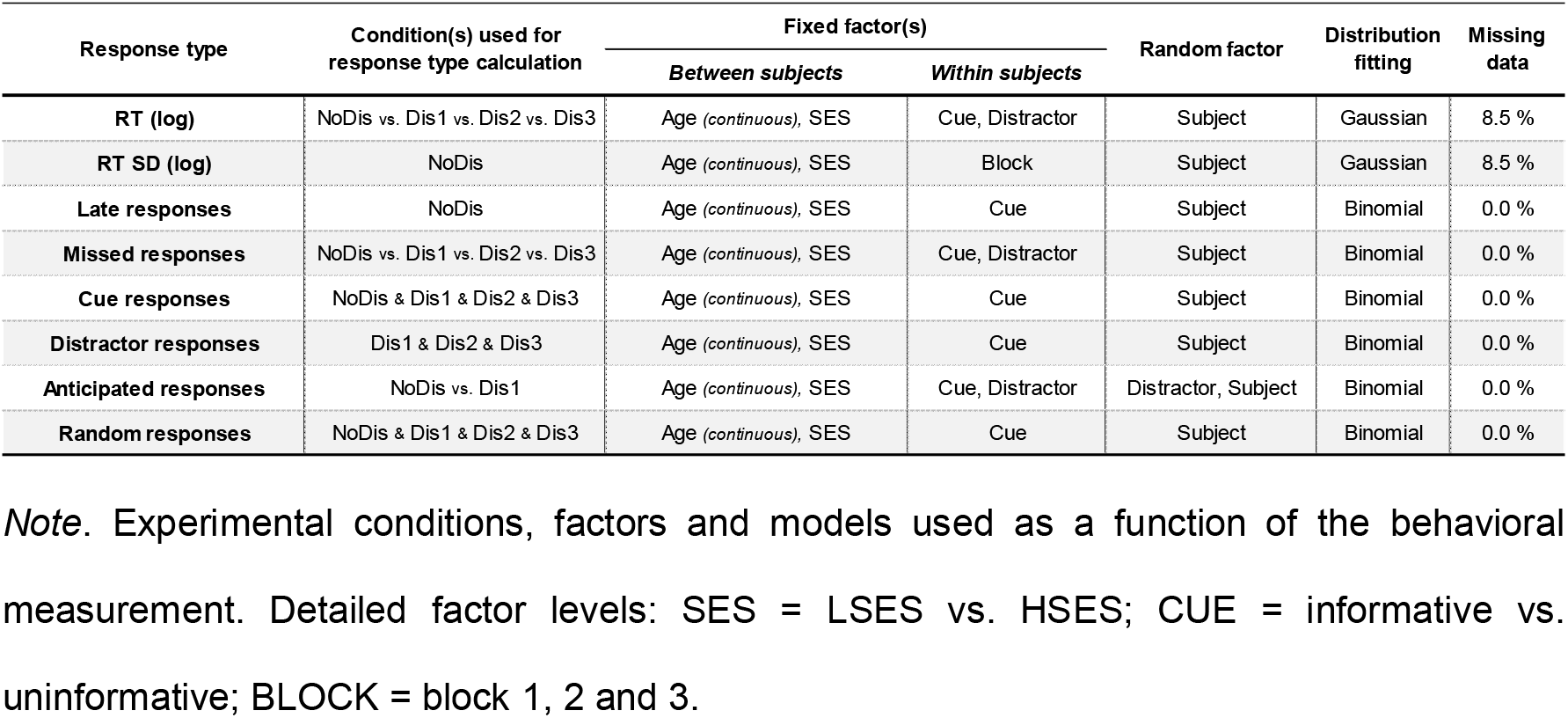
Main statistical analyses according to response types.

Frequentist models and statistics were performed in R, version 4.2.1, using the lme4 (Bates et al., 2015) and car (Fox & Weisberg, 2018) packages. Because both fixed and random factors were taken into account in statistical modelling, we ran a Type II analysis of variance. Wald chi-square tests were used for fixed effects in linear mixed-effects models (Fox & Weisberg, 2018). We only considered the explanatory variables. The fixed effect represents the mean effect across all subjects after correction for variability.

Wald chi-square test for mixed models is assumed to be anti-conservative. To ensure the robustness of the results, only main and interactions effects with a *p* value < .01 are considered in the followings.

When we found a significant main effect or interaction of the SES, CUE and DISTRACTOR factors, post-hoc Honest Significant Difference (HSD) tests were performed using the R emmeans package. *P*-values were considered as significant at *p* < .05 and were adjusted for the number of comparisons performed. To calculate effect sizes we used the R effectsize package: for linear mixed models (Gaussian distribution, RT) we calculated the ratio between measures, computed the omega squared coefficient and interpreted it using Cohen’s guidelines (Cohen, 1992); for generalized linear models (binomial distribution, response types) we computed the log odds ratio and interpreted it using Cohen’s guidelines (Cohen, 1988).

To confirm or infirm the presence of potential AGE effects on attention abilities, planned Bayesian Pearson correlations with age were performed on specific RT measures of attention: the attention orienting effect (median RT Uninformative – median RT Informative), the distraction effect (median RT Dis3 – median RT Dis1) and the phasic arousal effect (median RT NoDis – median RT Dis1). Bayesian statistics were performed using JASP® software (JASP Team (2018), JASP (Version 0.9) [Computer software]). We employed Bayes Factor (BF_10_) as a relative measure of evidence in favor of the null model (BF between 0.33 and 1, weak evidence; 0.1 to 0.33, positive evidence; a BF 0.01 to 0.1 strong evidence; BF lower than 0.01, decisive evidence) and the strength of evidence against the null model (BF of 1 to 3, weak evidence; BF 3 to 10, positive evidence; BF 10 to 100, strong evidence; BF higher than 100, decisive evidence; Lee & Wagenmakers, 2013).

To ensure that analyses were performed on a sufficient number of trials per condition, participants with fewer than 12 trials with positive RT in each of the distractor conditions (N = 6, all LSES) were excluded from the median RT analysis (resulting in a total average of trials with positive RT of 49 ± 1.8 in NoDis, 16 ± 0.7 in Dis1, 15 ± 0.6 in Dis2 and 16 ± 0.6 in Dis3 conditions across the overall sample). The revised sample size for median RT analysis was N=65. The percentage of missing data over the total sample of included subjects in analyses is shown in Table 3. Raw RT were log-transformed at the single trial scale for RT and RT SD analyses to be better fit to a linear model with Gaussian family distribution; response types were re-fit to a linear model with binomial distribution without transformation (see Table 3 and Appendix F for additional details).

In the Results section, we report the SEM as the estimator of the distribution dispersion of the measures of interest, when not specified.

## Results

### Data

For each type of behavioral measurement, we analyzed the influence of AGE (continuous), SES, CUE, and DISTRACTOR factors (unless specified otherwise in the Table 3). In the following, we only interpreted main and interaction effects with a *p* value < .01 to ensure the robustness of the results (i.e., to reduce Type I error). The results of the main analysis are summarized in table 4.

**Table 4.**
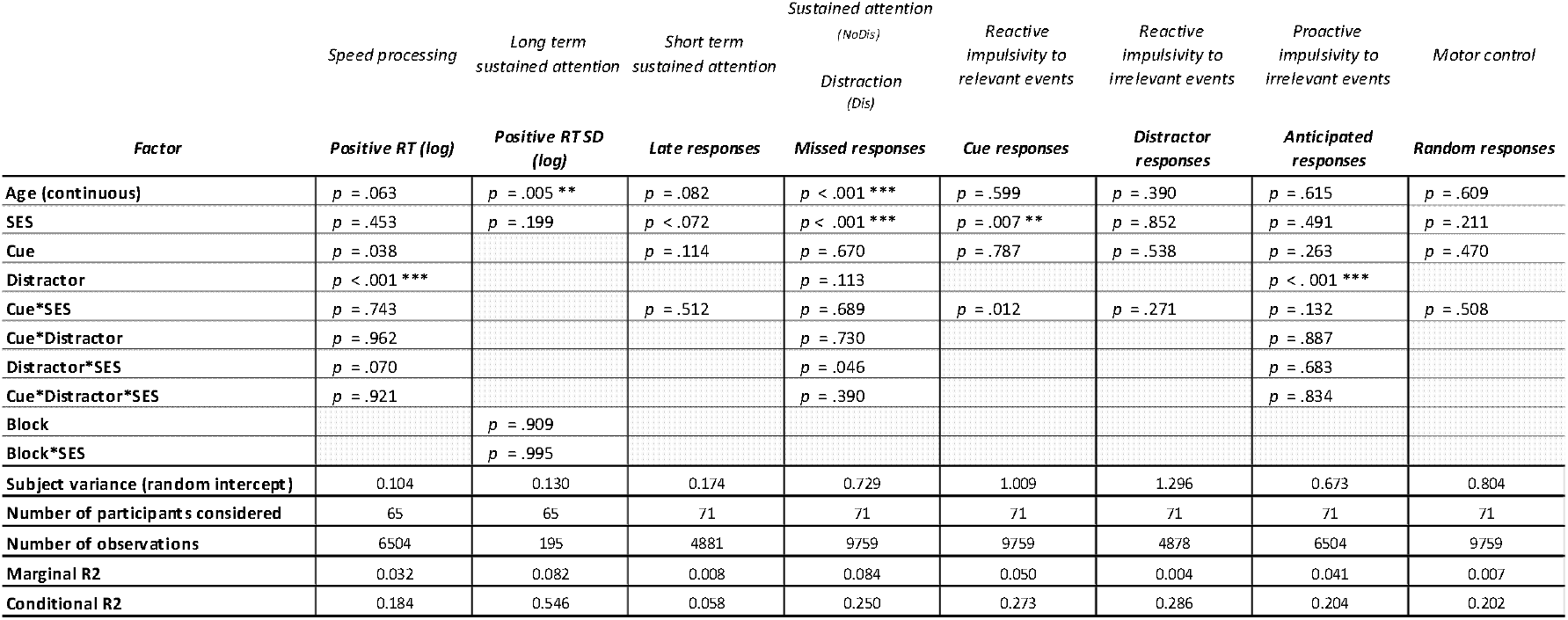
Results of the main statistical analyses, subject variance, number of participants, observations considered, as well as marginal and conditional R^2^ as a function of response type.

### Reaction Times

Consistent with our previous work in older children and adults, we observed a main effect of the DISTRACTOR (χ2 (3) = 180.4; *p* < .001; see Fig. 5) on RTs. Post-hoc pairwise comparisons showed that distractors speeded the response to targets. RTs were slower in NoDis (898.5 ± 35.0 ms) than in Dis1 (766.1 ± 39.0 ms, ratio NoDis / Dis1 = 1.36, CI [1.25, 1.46], effect size: large), Dis2 (801.5 ± 49.2 ms, ratio NoDis / Dis2 = 1.28, CI [1.18, 1.38], effect size: large) and Dis3 (848.2 ± 42.6 ms; ratio NoDis / Dis3 = 1.14, CI [1.05, 1.23], effect size: large) conditions. As shown in Figure 4, slower RTs were observed in Dis3 compared to Dis1 (ratio Dis3 / Dis1 = 1.19, CI [1.10, 1.28], effect size: large) and Dis2 (ratio Dis3 / Dis2 = 1.12, CI [1.04, 1.21], effect size: small) conditions, whereas no difference was found between Dis1 and Dis2 conditions (*p* = .345).

**Figure 4.**
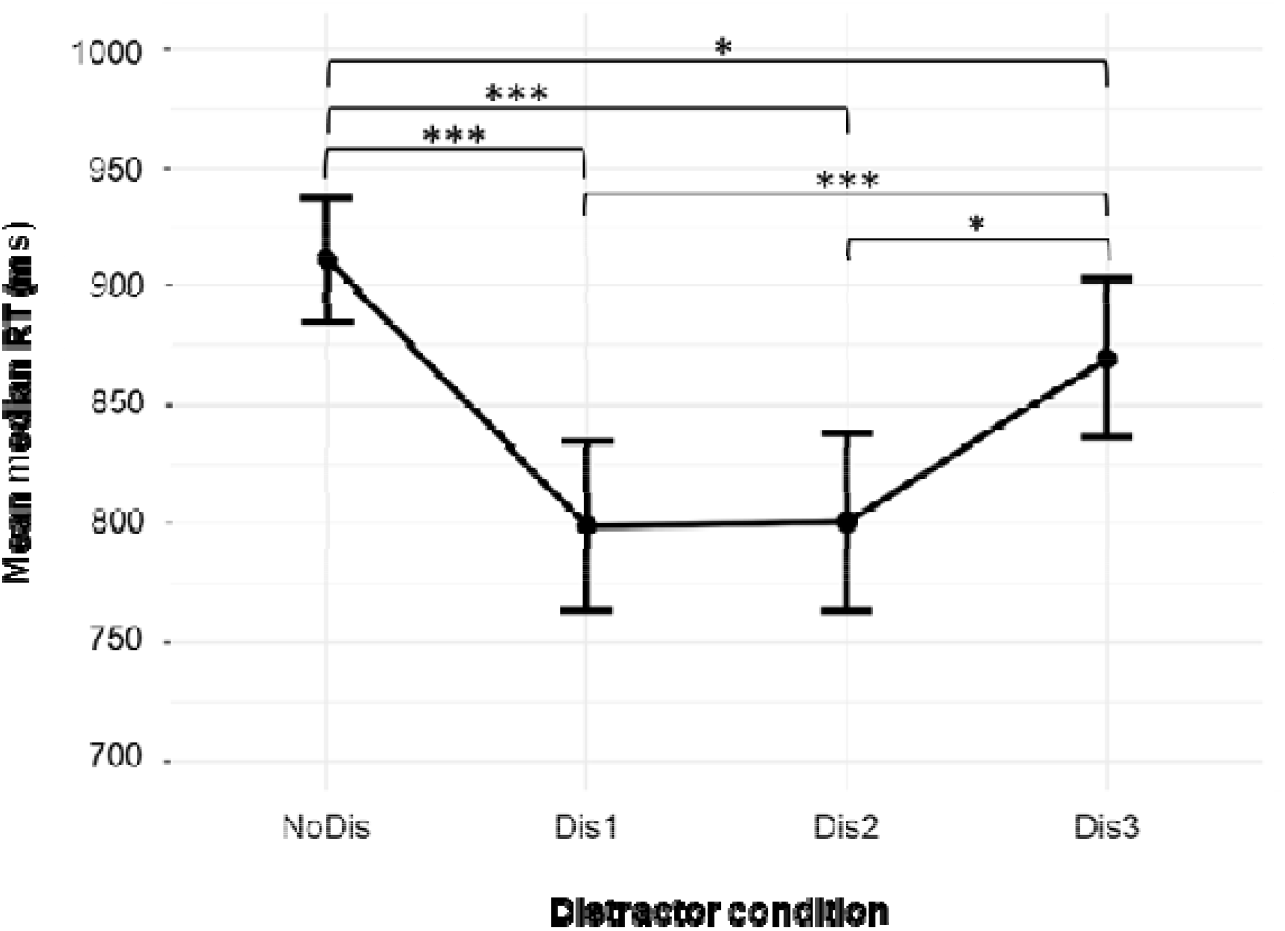
Median RT as a function of distractor condition. *Note*. Mean of median reaction time as a function of the distractor condition [NoDis, Dis1, Dis2 and Dis3] (*p* < .05 *, p < .001 ***; Error bars represent 1 SEM).

**Figure 5.**
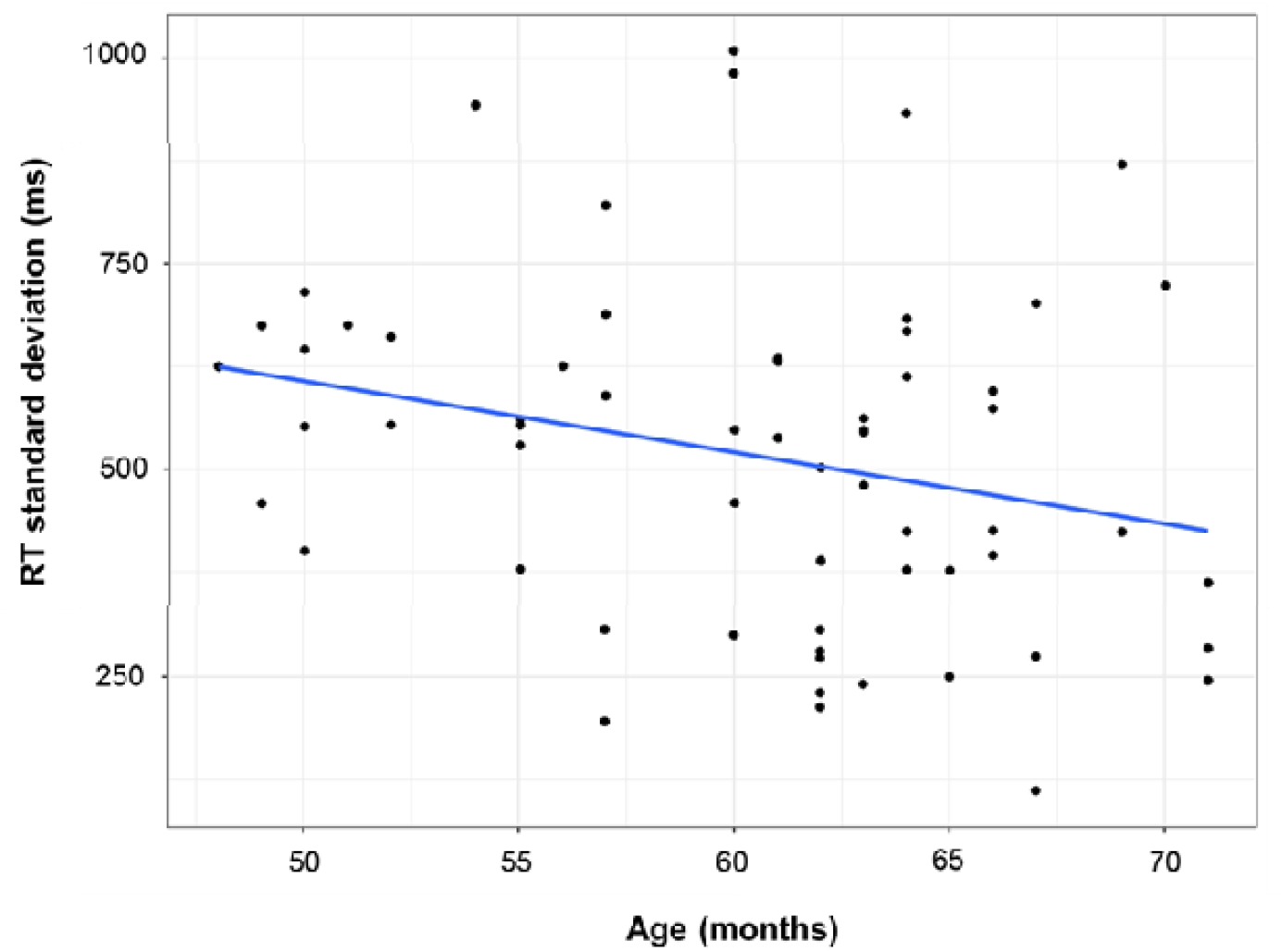
RT standard deviation as a function of age. *Note*. Mean RT standard deviation across blocks (1^st^, 2^nd^ and 3^rd^) as a function of age in months. Each dot represents the mean for a single subject.

No significant main effect of AGE was observed, suggesting no difference in speed of processing in children as a function of age.

Bayesian correlations were performed to investigate the effect of AGE on different distractibility components indexed by RT, namely the orienting, distraction and phasic arousal effects. The absence of an AGE effect on these different components was confirmed by Bayesian statistics (orienting measure and AGE: Pearson’s *r* = −0.12, BF_10_ = 0.24; distraction measure and AGE: Pearson’s *r* = −0.01, BF_10_ = 0.16; phasic arousal measure: Pearson’s *r* = −0.18, BF_10_ = 0.16). Overall, results from these analyses reveal positive evidence in favor of the null model: an absence of correlation between the given measures and the AGE.

### Standard deviation of reaction times

We observed a main effect of AGE (χ2 (1) = 5.5; *p* < .01; see Fig. 5) on RT standard deviation: this measure decreases with age, suggesting an increase in sustained attention from 4 to 5YO.

### Global accuracy

The proportion of different types of responses according to age is shown in Fig. 6. The average correct response rate was 50.7 ± 2.2%. No main effect of AGE or SES, nor interaction with SES, was found for random responses (total average: 6.0 ± 0.7%), distractor responses (total average: 5.9 ± 0.9%), anticipated responses (total average: 13.8 ± 1.1%) or late responses (total average: 9.9 ± 0.6%). Significant effects of AGE and SES on the other response types are detailed in the following sections.

**Figure 6.**
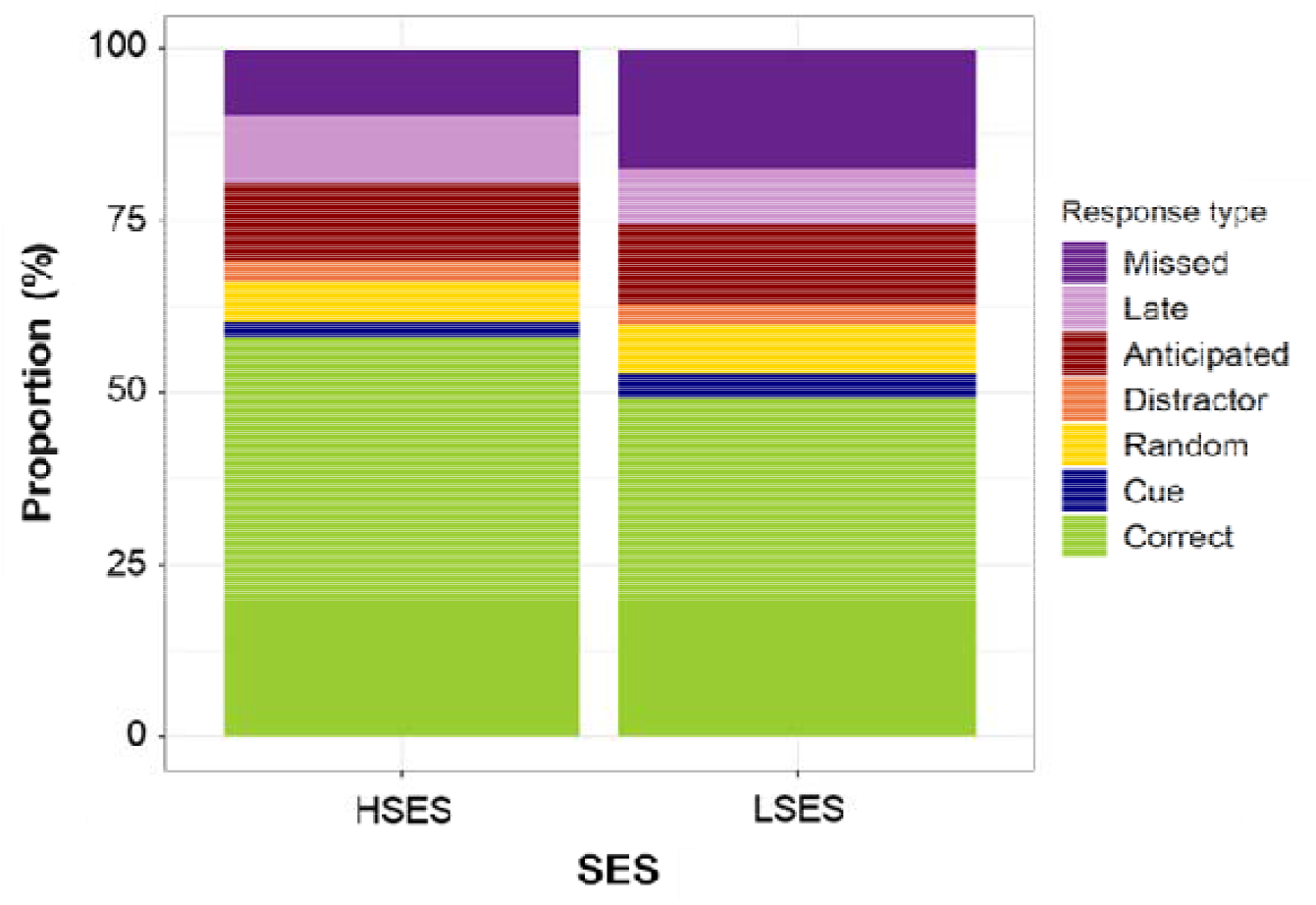
Response types proportions as a function of SES. *Note*. Response type proportions for LSES and HSES groups (HSES = higher SES, LSES = Lower SES).

### Erroneous responses to cue

The rate of cue responses (2.9 ± 0.4% on average) was modulated by SES (χ2 (1) = 7.0; p < .01; see Fig. 6): LSES children (3.7 ± 0.5%) made more cue responses than HSES children (1.6 ± 0.5%, log odds ratio LSES – HSES = 3.02, CI [1.59, 5.75], effect size: medium) irrespective of the validity content of the cue.

### Erroneous responses in anticipation of target

The rate of anticipated responses (13.8 ± 1.1% on average) was modulated by DISTRACTOR (χ2 (1) = 46.0; *p* < .001). Children had higher proportions of anticipated responses in Dis1 (18.7 ± 1.8%) than in NoDis condition (8.9 ± 0.9%, log odds ratio NoDis / Dis1 = 0.40, CI [0.33, 0.48], effect size: small).

### Missed responses

The rate of missed responses (15.1 ± 1.5% on average) was modulated by AGE (χ2 (1) = 15.4; *p* < .001; see Fig. 7): missed responses decreased with age. We also observed a significant main effect of SES (χ2 (1) = 15.0; *p* < .001; see Fig. 6). Children from LSES backgrounds (9.6 ± 1.6%) had more missed responses than their peers from HSES backgrounds (18.6 ± 1.2%, log odds ratio LSES – HSES = 2.46, CI [1.56, 3.88], effect size: small).

**Figure 7.**
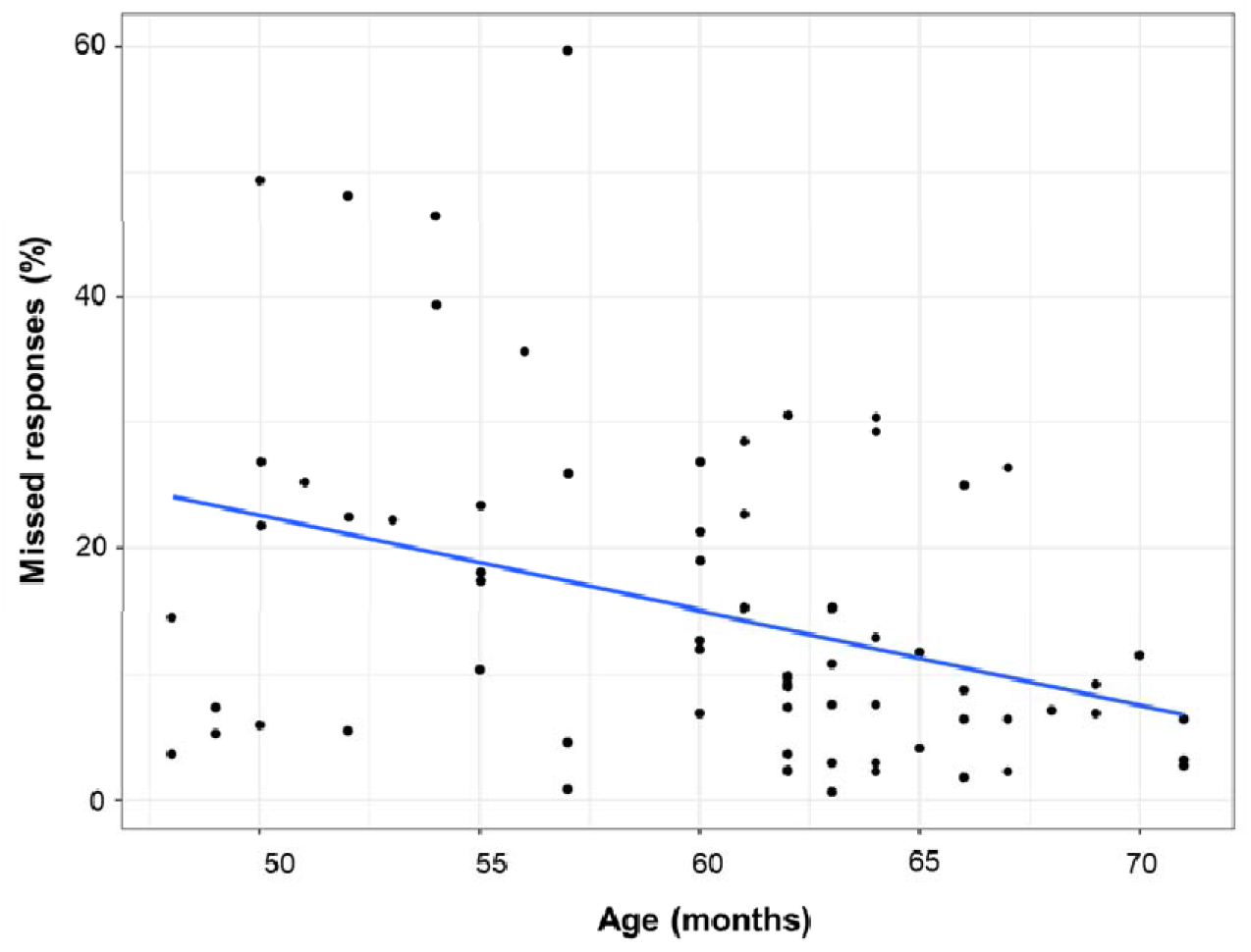
Missed responses as a function of age. *Note*. Missed response percentage as a function of the age in months. Each dot represents the mean for a single subject.

## Discussion

We characterized different components of distractibility in young children to determine the extent to which SES is related to the development of these components in early childhood. Consistent with our previous work in older children and adults, we found in 4- and 5-year-old children (i) a facilitation of RT after early distractors, (ii) a cost in RT after late distractors (iii) a high variability in RT, suggesting that this paradigm provides valid measures of these constructs in younger children (see also Appendix G, Fig. G1, for a visual representation of the developmental trajectories of distractibility components from age 4 to 25). Furthermore, LSES children made more erroneous responses to the cue and missed more targets than children from the HSES group, suggesting greater impulsivity and reduced sustained attention in the LSES group. Below, we discuss the implications of these findings, caveats to our conclusions, and future directions.

### Difference in distractibility components from 4 to 5 years of age

#### Voluntary orienting and sustained attention

To avoid Type I error, the alpha threshold to detect a factor effect on the different CAT measures was set at *p* < .01. According to this threshold, we did not observe a significant effect of the cue on RT, thus indicating that both 4 and 5YO, independent of SES, were equally fast to respond after informative and uninformative cues. This result suggests that voluntary attention orienting is still immature during the preschool period. In a previous CAT study (Hoyer et al., 2021), we observed an adult-like voluntary attention orienting effect in school-aged children (6-17 years old). As previously suggested, it is possible that attention is still maturing in preschoolers before reaching an adult-like efficiency at age 6 (Hrabok et al., 2007; Mezzacappa, 2004; Posner et al., 2014; Rothbart et al., 2011). But voluntary attention orienting may also be highly variable in young children, thus making this effect more difficult to detect in preschool-aged children; this might explain why some studies also found similar voluntary attention orienting effects in preschoolers and adults (Colombo, 2001; Johnson et al., 1991; Ross-Sheehy et al., 2015). Further studies are needed to better understand inter-individual differences in voluntary attention orienting during the preschool age.

Reaction time standard deviation has been found decreasing from 4 to 5 years of age, suggesting that sustained attention improves during the preschool period: this result is consistent with previous findings (Graziano et al., 2011; Reynolds & Romano, 2016; Richards & Casey, 1991; Ruff & Capozzoli, 2003). Furthermore, the number of missed responses in attentional tasks is a sensitive measure of sustained attention over long time periods (Kanaka et al., 2008), while the proportion of late responses is considered to be an index of short-term sustained attention (Petton et al., 2018). In the present study, and irrespective of the distractor condition, the missed response rate decreased with age suggesting that sustained attention abilities are still developing during the preschool years. However, no effect of age was observed for the late response type, suggesting that children’s ability to maintain attention over shorter periods of time is more stable from 4 to 5 years of age. Increased missed responses in younger children are then more likely to result from an overall lower vigilance level in the younger age group, though future research is necessary to confirm this hypothesis. In addition, all tested 3YO children were excluded from analysis due to their random performance; an inability to adequately perform the task at 3 years of age could also be explained by reduced sustained attention abilities, as previously observed (Mahone et al., 2001). Children whose performance was excluded from the analysis were not able to stay focused on the task according to the instructions (e.g., looking at the screen, staying sit, not talking too much during the test); their performance thus likely does not reflect attention efficacy as it is intended to be measured by the CAT. Among these excluded children, a majority came form LSES backgrounds: this phenomenon may reflect a developmental lag in attention ability in this population, which is consistent with ERP evidence from preschool-aged children from different SES backgrounds (Hampton Wray et al., 2017). The CAT is relatively demanding because participants need to maintain focus to answer frequent targets, suggesting that the paradigm might not be appropriate to measure distractibility if attention is not sufficiently developed. Further studies that seek to better understand interindividual differences in attention are needed to precisely identify which child characteristics can predict their capacity to perform the CAT.

Together, these results suggest that sustained attention as measured by the CAT is still developing between ages 4 and 5, but may not be sufficiently developed at age 3 and in some four-year-old children (particularly those from LSES backgrounds) to obtain reliable measures.

#### Distraction and facilitation effects triggered by distractors

In line with previous studies using the CAT in adults (Bidet-Caulet et al., 2015; ElShafei et al., 2019, 2020; Masson & Bidet-Caulet, 2019), we observed two distinct effects on RT triggered by the distracting sounds. First, distracting sounds played long before the target (Dis1 and Dis2) speeded RT compared to a condition without distractor (NoDis): this benefit in RT has been attributed to an increase in phasic arousal (Masson & Bidet-Caulet, 2019). Second, distracting sounds played just before the target (Dis3) slowed RT compared to conditions with a distractor played earlier (Dis1 and Dis2); this cost in RT is considered a good behavioral approximation of distraction (Bidet-Caulet et al., 2015; ElShafei et al., 2019; Hoyer et al., 2021; Masson & Bidet-Caulet, 2019). In addition, both phasic arousal and distraction effects were observed in 4-5-year-olds. Thus, this finding extends those previously obtained with school-aged children (Hoyer et al., 2021) to preschool-aged children.

To date, few studies have investigated distraction in preschool-aged children using active behavioral tasks. Recent results from oddball paradigms, however, suggest that the distraction effect decreases from age 4 to 6 and 6 to 10 before reaching the adult level (Wetzel et al., 2018). On the contrary, in the present study, the RT distraction effect is similar between 4- and 5-year-olds. This discrepancy may be explained by differences in protocols. Discrimination (Oddball task; Wetzel et al., 2018) is more demanding than detection (CAT): it could then be easier for 4YO children to deal with distracting information during detection tasks compared to discrimination tasks. Thus, future studies using both different task difficulty levels in the same sample of children are necessary to improve our understanding of the development of distraction during the preschool period.

#### Motor control and impulsivity

In the CAT paradigm, anticipated, cue, distractor and random responses are conceptualized as complementary measures of impulsivity and motor control. The present findings indicate that none of these measures were modulated by age, suggesting that impulsivity and motor control efficiency are not different between 4 and 5 years of age. Impulsivity has previously been found reducing during the early preschool period (i.e., between age 3 and 4, Wiebe et al., 2012), and then progressively decreases from the preschool age to early adulthood (Cragg & Nation, 2008; Hoyer et al., 2021). Furthermore, irrespective of age, participants made a few more anticipated responses to the target in presence of a distractor, showing that processes triggered by distractors contribute to increase impulsivity to subsequent targets in preschool-age children.

### Differences in distractibility as a function of SES

#### Distraction

No effect of SES was found on any of the distraction measures provided by the CAT: enhanced RT, late response or missed responses in trials with distracting sounds. Therefore, the present study does not provide clear support for increased distraction in children from LSES backgrounds.

This finding contradicts previous research seeking to characterize behavioral differences in attention during the preschool years and as a function of SES. Results from event-related potential (ERP) studies of auditory selective attention suggested a reduced ability to suppress distraction in LSES children (Stevens et al., 2009; Giuliano et al., 2018; Hampton Wray et al., 2017). This discrepancy may be explained, at least in part, by differences in the tasks used to assess attentional capacities. Studies suggesting a reduced ability to suppress distracting information in LSES children have used a dichotic listening task with embedded probes. Behaviorally, this paradigm solicits the child’s ability to inhibit the continuous stream of distraction from the unattended channel, but also unexpected and isolated probe sounds. Since these tasks are more demanding, a child may need to recruit the executive system, the development of which is delayed in children from LSES backgrounds (Lawson et al., 2013; Noble et al., 2012). Recent findings also suggest that for children from LSES backgrounds there may be a potential biological cost of achieving performance similar to their peers from HSES backgrounds (Giuliano et al., 2018). Future studies that combine different paradigms in the same sample of LSES and HSES children are necessary to further elucidate this pattern of results.

#### Sustained attention

Although behavioral distraction was similar for children from LSES and HSES backgrounds, we found evidence that 4 and 5-year-old children from LSES backgrounds had more difficulty in sustaining attention, as they missed more targets irrespective of the distractor condition. This difference as a function SES, while a small effect, is consistent with evidence from other studies. Previous studies using a child-friendly ERP paradigm assessing sustained selective auditory attention (i.e., the ability to focus on a relevant channel among others over time) have also found differences in brain activity as a function of SES in this age group (Stevens et al., 2009; Hampton-Wray et al., 2017, Giuliano et al., 2018). There is also evidence that preschool-aged children from LSES backgrounds show differential functional maturation of prefrontal cortices (Lawson et al., 2013; Noble et al., 2012). Taken together, these results suggest that preschool-aged children from LSES backgrounds may show an attentional imbalance characterized by reduced voluntary sustained attention. This might represent an adaptation to certain environmental contexts in which regularly interrupting ongoing voluntary attention processes is necessary to maintain constant reactivity to possible changes in the environment.

#### Motor control and impulsivity

Children from LSES backgrounds made significantly more cue responses than their peers from HSES backgrounds, suggesting higher impulsivity in LSES children in line with previous findings (Arán-Filippetti & Richaud de Minzi, 2012; Ruf et al., 2008). Importantly, this present finding further suggests that differences in impulsivity as a function of SES may be specific to relevant stimuli (responses to cues), as impulsivity to these stimuli was increased in children from LSES backgrounds, while there was no difference in SES groups in impulsivity to irrelevant stimuli (responses to distractors).

#### Limitations of the present study

There are several limitations to the current study. First, the protocol used does not provide measures of stress, motivation, or tonic arousal, all of which are believed to influence attentional performance. Second, this cross-sectional study did not provide longitudinal data. Further studies are then needed to characterize individual variability in attention performance and to better identify predictors of attentional efficiency in children and examine developmental trajectories of efficiency in the same participants. Third, SES was operationalized by proxy using the school affiliation of the child, limiting the ability to characterize and control for variability within SES. The present results thus need to be replicated using a more rigorous, individual-level operationalization of SES that characterizes multiple dimensions of the construct (Hackman et al., 2010; Ursache & Noble, 2016), and with larger samples. It is also important to note that SES is not the sole factor that can explain differences in attention during the preschool period; several other related variables, such as the time spent by parents playing with their child (e.g., Cheung et al., 2017) or the time spent by children participating to activities outside the class (e.g., Diamond et al., 2007) have been shown to have a positive relationship with child cognitive abilities. It will be fruitful for future studies attempting to replicate the present results to include such factors in their design and analyses. Finally, a substantial large number of children were excluded because they did not manage to perform the task according to the instructions (e.g., look at the screen, stay seated and quiet during the test); therefore, results from this study may not be generalizable to a minority of children, whose attention would not be sufficiently developed enough due to individual differences in development. It will thus be important to assess distractibility in larger samples of children, and to employ different attention tasks to better characterize inter-individual variability in attention performance within SES groups (e.g., 16 LSES and 5 HSES children were excluded from analyses because the task was too difficult).

#### Conclusion and future directions

To our knowledge, this is the first study to explore differences in specific facets of distractibility during the preschool period. The attention task we used, the CAT, was found to be suitable for a majority of 4- and 5-year-old children, but a minority of these children (particularly from LSES backgrounds) and 3-year-old children were unable to complete the task according to the given instructions. This suggests that individual differences in distractibility and attention are present during the preschool developmental period, and that the CAT is not suitable for every preschooler. Nevertheless, the present study permitted the characterization of distractibility in children who had sufficient voluntary sustained attention ability to perform the task according to the instructions. Results show that voluntary sustained attention ability increases from 4 to 5 years of age as indexed by increased reaction time variability and a greater number of missed responses in the younger children. Sustaining attention over time was found to be more difficult for children from LSES backgrounds, but no evidence for enhanced distraction in this population was found. While this difference in sustained attention may be maladaptive in certain contexts like school, it is important to consider that regularly interrupting ongoing attentional processes may be adaptive by allowing a child to maintain more constant reactivity to possible changes in the environment. Elucidating differences in distractibility during the preschool period as a function of SES is relevant not only to our basic understanding of disparities related to SES, but also for the timing or nature of interventions targeting attention skills (Neville et al., 2013; Posner et al., 2015; Tang & Posner, 2009). Thus, this study showed that the CAT is a potentially powerful tool for assessing attention in preschool-aged children whose sustained attention is sufficient to perform the entire task, and that the CAT can provide important information for further evidence-based approaches to develop and refine programs that seek to train attention processes in children.

## Notes

### Competing Interest Statement

The authors have declared no competing interest.

### Summary of Updates

Clarifications in the method section and effect sizes added.

